# Breaking the sparsity barrier in clinical targeted-panel sequencing: Mapping the inherited determinants of mutational signatures

**DOI:** 10.64898/2026.03.29.714525

**Authors:** Alon Ravid, Hagay Ladany, Alexander Gusev, Yosef E. Maruvka

**Affiliations:** Department of Mathematics, Technion – Israeli Institute of Technology, Haifa, Israel; Department of Biotechnology and Food Engineering, Technion – Israeli Institute of Technology, Haifa, Israel; Department of Medical Oncology, Dana-Farber Cancer Institute & Harvard Medical School, Boston, MA, USA; Program in Medical and Population Genetics, Broad Institute, Cambridge, MA, USA

## Abstract

Cancer development is shaped by somatic mutational processes that leave characteristic patterns known as mutational signatures. The inherited determinants of variability in signature activity remain largely unknown. Common germline variants that regulate this activity, which we term Signature Quantitative Trait Loci (SigQTLs), are expected to have modest individual effects, requiring cohorts of tens of thousands of samples for reliable detection. Clinical targeted-panel sequencing datasets achieve this scale, but present a fundamental challenge: individual tumors typically harbor too few mutations for stable signature inference. To overcome this sparsity barrier, we introduce GroupSig, a framework that aggregates sparse mutational patterns across samples sharing a germline genotype into information-rich meta-samples, enabling robust signature inference at the population level.

We validated GroupSig by recovering the well-established correlations between age and clock-like signatures SBS1 and SBS5 using emulated panel data from The-Cancer-Genome-Atlas. We then applied GroupSig to approximately 32,000 tumor samples from the Dana-Farber Cancer Institute PROFILE cohort in a genome-wide SigQTL scan. We identified 9 genome-wide significant SigQTLs, with the strongest signal at locus 16q24.3, where six variants were associated with increased SBS7 (UV exposure) activity. This association persisted after excluding melanoma samples, arguing against a tumor-type enrichment artifact. Validation in TCGA confirmed 6 SigQTLs, all at 16q24.3, where implicated variants are eQTLs for CDK10 and SPG7 in skin tissue. Beyond genome-wide hits, DNA repair genes were 12.6-fold enriched among sub-threshold signals, supporting a polygenic architecture for mutational process regulation. GroupSig provides a scalable framework for germline-somatic association studies using panel sequencing data.

## Introduction

Cancer development is driven by the accumulation of somatic mutations, resulting from stochastic events and various mutational processes^1^. Certain mutational processes (e.g., DNA mismatch-repair deficiency, UV and tobacco exposure) generate mutations in specific trinucleotide contexts also known as mutational signatures^2^. These signatures may help infer the underlying mechanisms of tumorigenesis and cancer evolution and inform prevention and therapy^3^. In a given sample, multiple mutational processes can imprint distinct mutational signatures, producing the sample’s composite mutational pattern that obscures the process of origin of individual mutations. The mutational pattern of a sample can be deconvolved, e.g., using non-negative matrix factorization (NMF), to infer the activity of each signature^4,5^.

Signature activities vary across tumors and patients^2,6–8^, and this variability is not purely stochastic: inherited and environmental^9–11^ factors can influence the activity of mutational processes. The determinants of this variability remain incompletely understood, particularly in tumors without clear exogenous exposures or canonical DNA-repair defects. One possible mechanism is that common germline variants can subtly influence DNA repair efficiency or resistance to DNA damage.

Genome-wide association studies (GWAS) have linked common germline variants to numerous human phenotypes, including cancer risk^12^. In addition to increased cancer risk, germline variants or loci may also influence somatic mutational processes - loci we term Signature Quantitative Trait Loci (SigQTLs). Analogous to expression QTLs (eQTLs)^13,14^, which quantify the effect of germline variants on gene expression levels, SigQTLs quantify their effect on the activity of somatic mutational processes. However, whether and how common inherited variants modulate somatic mutational processes remains largely underexplored^8^. Detecting the effect of common germline variants requires extensive cohorts (>10,000 samples) because the effect of individual variants is expected to be modest. Current whole-genome sequencing (WGS) and whole-exome sequencing (WES) datasets are limited in size, rendering them unfeasible for such analyses. On the other hand, clinical targeted-panel sequencing cohorts possess the necessary sample volume but fail to provide the number of mutations per sample needed for inferring signatures activity which is typically around 30 or so^15^. In targeted panels, the mutational pattern is too sparse, thus signature activity is often not reliably estimable per sample, preventing standard per-sample association testing.

To address this limitation, we developed GroupSig (Fig. 1), a framework that aggregates somatic single base substitutions (SBS) across patients sharing a specific feature (e.g., a germline variant, age, or tumor subtype) into ‘meta-samples’. This aggregation generates a meta-sample with a dense enough mutational pattern that has the sufficient number of mutations required for signature inference, effectively overcoming the data sparsity an individual sample has. Consequently, GroupSig facilitates association testing between mutational signature activity and a given feature. We validated GroupSig by testing its ability to infer known age association of age-related signatures. We then applied GroupSig to a targeted panel sequencing cohort (∼32,000 tumors) from the Dana-Farber Cancer Institute (DFCI) to perform a genome-wide scan for SigQTLs, and validated significant associations in the TCGA dataset.

**Figure 1.**
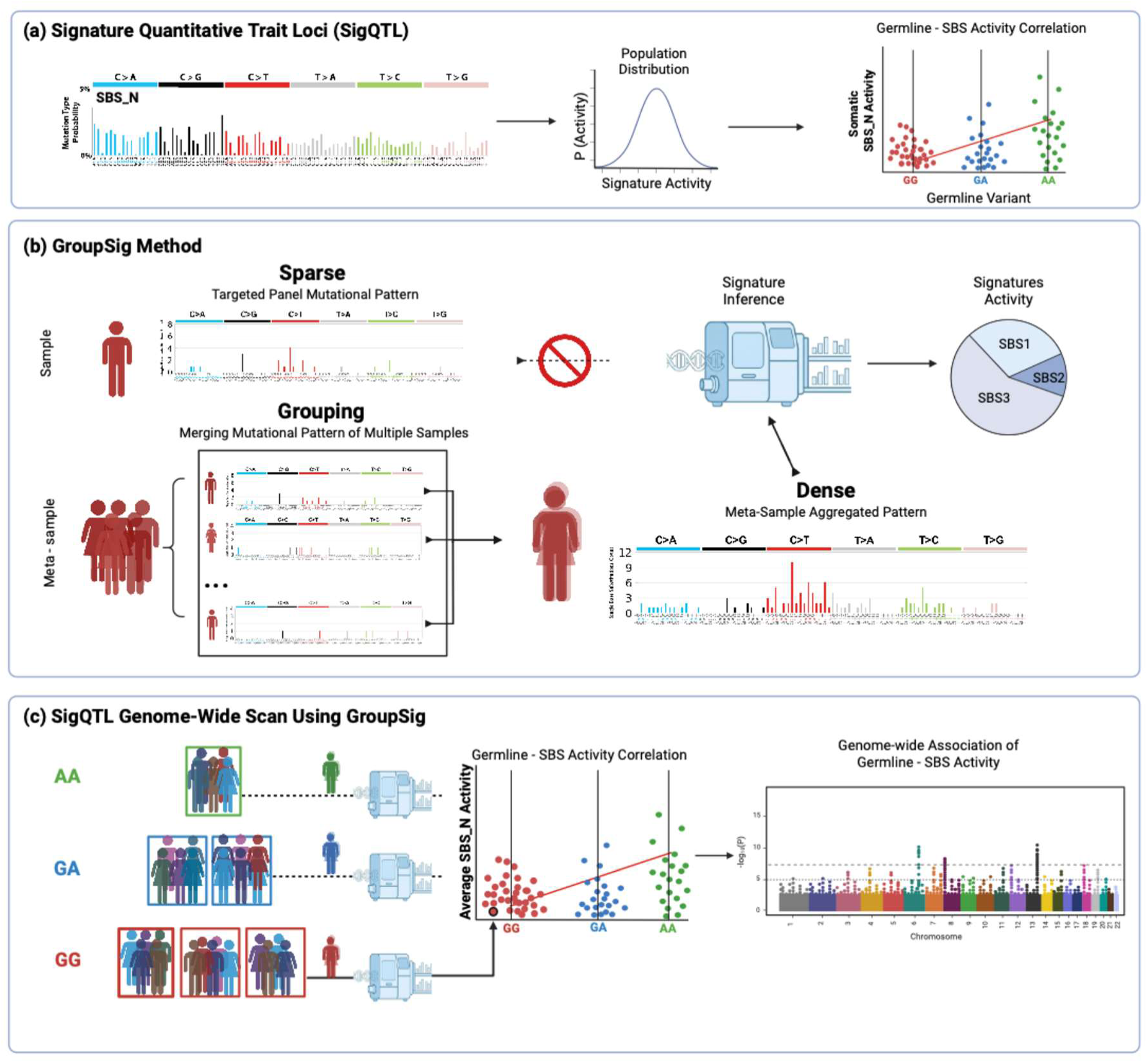
Schematic description. **(a)** Signature Quantitative Trait Loci (SigQTL). The activity of a mutational signature (e.g., SBS_N) varies across individuals depending on their germline genotype at a specific variant. Differences in genotype (e.g., GG, GA, AA) lead to distinct distributions of signature activity, which can be tested through correlation between the genotypes and somatic SBS signature activity. **(b)** GroupSig Method. Targeted panel sequencing typically produces sparse mutational profiles for individual samples, preventing reliable mutational signature inference. To overcome this limitation, we introduce GroupSig, which aggregates mutational patterns from multiple samples sharing a common feature (e.g., genotype) to generate meta-samples. By combining patterns across samples, the resulting aggregated profile becomes sufficiently dense to enable accurate mutational signature inference. **(c)** SigQTL Genome-wide Scan Using GroupSig. For each germline variant, samples were stratified by genotype (e.g., AA, GA, GG) and, within each genotype class, randomly partitioned into disjoint groups of equal size. Mutation patterns were aggregated within each group to create meta-samples, and signatures activity for each meta-sample were inferred using MuSiCal (supervised refitting to known SBS profiles). Average activity of each signature per meta-sample was modeled by linear regression. Variants exceeding the genome-wide significance threshold (p < 5 × 10−8) were annotated as SigQTLs.

## Materials and methods

### Data and preprocessing

We analyzed two datasets: (i) The DFCI Profile clinical targeted-panel cohort that was applied to ∼32,000 tumors, which was used as the discovery cohort. (ii) The TCGA whole-exome sequencing cohort (∼10,000 tumors), which was used for method development and as a validation set. All analyses were performed under institutional approvals and data-use agreements.

DFCI tumor specimens (n = 31,984) were obtained from patients at the DFCI who provided written informed consent for the Profile prospective clinical sequencing effort (IRB protocol 11-104). Pathologists reviewed all specimens to ensure a minimum tumor cellularity of 20%. Secondary analyses of this dataset were performed under DFCI IRB protocols 19-033 and 19-025 with a waiver of HIPAA authorization. Samples were sequenced using one of three versions of the OncoPanel targeted capture platform, covering 275, 300, and 447 genes, respectively. Technical requirements included a minimum of 30× coverage for 80% of targets, with average unique depths of approximately 200× for version 1 and 350× for versions 2&3. Reads were aligned using BWA^16^, flagged for duplicates with Picard Tools, and locally realigned via GATK^17^. SBS were called using MuTect, followed by manual review. To isolate somatic alterations, putative germline variants were filtered out if present in a panel of historical normal or the Exome Sequencing Project (ESP) at a frequency ≥0.1%, unless the variant was documented in COSMIC.

TCGA mutations and batch-normalized gene expression levels were generated by the MC3 consortium^18^ as the consensus of different tools, and were obtained from the cbioportal website^19^.

#### Imputed germline

As we described elsewhere^20^, we performed DFCI germline imputation across all samples using STITCH; the targeted sequencing panels generated ultralow genome-wide data which can be leveraged for germline imputation^21,22^. We restricted germline variants to those with imputation INFO > 0.4 and minor allele frequency > 0.01. We then computed continental ancestry using imputed dosages and the PLINK2^23^ ‘--score’ function by projecting each sample into the reference principal component (PC) space generated by SNPweights tools in HapMap populations of European, West African (Yoruban), and East Asian (Chinese) ancestry^24^. We calculated the mean and standard deviation of PC1 and PC2 for self-reported white individuals and retain all individuals within two standard deviations of the mean (PC1: 1.58 × 10^−8^ [±1.05 × 10^−9^], PC2: −8.14 × 10^−9^ [±2.32 × 10^−9^]) in order to reduce population stratification biases^20^. After applying our sample and site filters (including reapplying the MAF filter to the final sample set after somatic mutation filtering described below), the working germline dataset contained n = 3,170,862 variants.

We performed TCGA genotype inference using raw Affymetrix SNP 6.0 CEL files using Affymetrix Power Tools (APT) v2.11.2^25^. We utilized apt-probeset-genotype binary with the BRLMM-P algorithm, applying the generic_prior.txt model and na35 annotation release. We converted the output CHP files to VCF format via bcftools v1.10.2 (plugin: affy2vcf)^26^, annotated using the NetAffx na35 manifest, and normalized against the GRCh37/hg19 reference. For imputation, the VCF files were then processed via the Michigan Imputation Server (v1.5.7)^27^. We executed phasing using Eagle v2.4^28^, and followed with imputation using Minimac4 v1.0.2, referencing the Haplotype Reference Consortium (HRC r1.1) panel for the European subpopulation^29^.

### The GroupSig method

GroupSig is a framework for inferring mutational signatures from aggregated rather than individual samples (Fig. 1). By pooling mutations from samples that share a defined characteristic, we construct “meta-samples” with substantially increased mutation counts, enabling robust signature inference from sparse panel datasets. The approach is agnostic to the specific inference algorithm.

The GroupSig workflow consists of four primary steps:

1. Partitioning: Samples are stratified based on the feature of interest.
2. Aggregation: Within each stratum, samples are partitioned into disjoint subgroups of size *n*, and their mutational patterns are summed to form a meta-sample mutational pattern. The group size n is calibrated to ensure that the number of cumulative mutations per meta-sample is sufficient for robust signature activity inference.
3. Inference: We then perform signature activity inference on the aggregated mutational patterns of the meta-samples.
4. Statistical analysis: The relationship between the feature and the inferred signature activity is statistically evaluated across meta-samples, while accounting for potential covariates such as ancestry, sex and more.

Hypermutated tumors can dominate aggregated meta-sample patterns and introduce bias. To reduce this risk and limit spurious associations, we excluded the top 10% of samples by somatic mutations count within each tumor type or samples with more than 40 mutations.

### Partition and Aggregation of mutations

For each tumor, we summarize somatic SBS mutations as a 96-trinucleotide pattern. Specifically, each sample is represented by a 96-dimensional count vector whose entries correspond to the six base-substitution classes stratified by the immediate 5′ and 3′ sequence context (e.g., a C>T substitution in 5’…ACG…3’ is encoded as a A[C>T]G trinucleotide context). Let *s_j_* denote the mutational pattern of sample *j*, and let *S* = {*s*_1_, …, *s*_N_} be the set of patterns across all samples. In standard supervised signature inference, these per-sample patterns are used directly to infer signature activity; however, in targeted-panel data they are typically too sparse for stable inference.

GroupSig increases mutational abundance by aggregating samples that share a feature of interest (e.g., germline genotype, tumor subtype, or a binned/ordered clinical variable). For a categorical feature with states *G* = {*g*_1_, …, *g*_L_}, we first stratify the cohort into subsets:

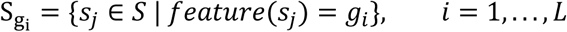

Within each stratum *g_i_*, samples are randomly partitioned into disjoint ‘meta-samples’ of size *n*. These meta-samples are constructed as non-overlapping sets; to handle remainders, the final group in each stratum incorporates any leftover samples, resulting in a terminal group size between *n* and 2*n* − 1. For each meta-sample *M_k,i_*, we compute an aggregated pattern by summing the constituent sample trinucleotide context pattern:

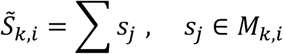

Each *S̃_k,i_* is therefore the meta-sample mutational pattern (i.e., its 96-dimensional trinucleotide contexts count vector) with substantially higher mutation counts than any individual sample, enabling robust signature inference.

For a continuous feature (e.g., age), we sort samples by the feature value and then form disjoint consecutive groups of samples of size *n*. The feature value assigned to each meta-sample is taken as the average of the feature among its member samples. The resulting aggregated patterns *S̃_k,i_* and their corresponding meta-sample feature values are then used as the inputs for downstream signature inference and association testing.

### Mutational signature inference

We inferred mutational signatures activity for a meta-sample using supervised signature inference methods. For a given single meta-sample *M*_k,i_,, its mutational pattern *S̃_k,i_* represents its aggregated mutational pattern (a 96 entries vector). This vector is projected on a reference matrix of *p* signatures *W* (96 × *p*) in order to compute a non-negative activity vector *A* (of size *p*) that satisfies:

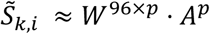

The resulting activity vector *A* contains the estimated number of mutations each signature generated in the observed composite pattern *S̃_k,i_*.

We evaluated several tools for signature activity inference: (i) SigProfiler^30^ (ii) SignatureAnalyzer^31^, packages which implement supervised non-negative matrix factorization^4,5^ (NMF) and (iii) MuSiCal^32^ which implements a likelihood-based non-negative least squares (NNLS). Given the computational magnitude of a genome-wide scan - requiring signature inference for thousands of meta-samples across ∼3.1 million loci - processing speed was the critical selection criterion. Consequently, we selected MuSiCal, which demonstrated the highest computational efficiency.

Because supervised signature inference can assign very small positive weights to a signature (overfitting/noise)^15^, we applied a simple post-inference filter to remove negligible contributions. For each meta-sample, we set to zero any signature whose activity accounted for less than 2% of those meta-sample’s total mutations. By implementing this 2% cutoff, we prioritize the removal of noise and overfitting over the retention of very low-activity signatures, which ultimately lack the necessary signal-to-noise ratio for robust correlation analysis.

### Signature selection

To minimize overfitting and improve inference stability, we used the more compact COSMIC v2 reference set (available at https://cancer.sanger.ac.uk/cosmic/signatures_v2). Although COSMIC v3 is more comprehensive, it also contains several signatures with high similarity, which increases the risk of misattribution during inference^33^.

To generate a stable, high-confidence signature inference, we excluded signatures that are (i) flagged as sequencing artefacts or as unvalidated^7,34^. (ii) present in fewer than 100 tumor samples or in fewer than three distinct tumor types in the TCGA dataset (that was used to generate the signatures repertoire) (iii) associated with hypermutation, mismatch repair deficiency or POLE deficiency, namely SBS6, SBS10, SBS14, SBS15, SBS20, SBS21 SBS26 and SBS28^7^. The exclusion of hypermutated related signatures is needed as we excluded samples that are highly mutated. After applying these criteria the final set comprised these signatures: SBS1, SBS2, SBS3, SBS4, SBS5, SBS7, SBS13, SBS17 and SBS18 (Table S1&2).

### Statistical analysis

To test associations between mutational signature activity and genotype, we used multivariable ordinary least squares (OLS) regression, as implemented in Python’s statsmodels.formula.api.ols function^35^. Genotypes were coded additively as 0, 1, or 2 for homozygous major, heterozygous, and homozygous minor genotypes, respectively. Models included sex (female = 0, male = 1), age, and the first 20 genetic ancestry principal components (PCs) as covariates. Variants were considered significant if the genotype coefficient in the OLS model differed from zero (two-sided test) after adjustment for covariates, using the conventional genome-wide significance threshold p < 5×10^−8^.

We computed PCs using PLINK 1.9^23^: we performed per-chromosome quality check and linkage disequilibrium (LD) pruning and excluded ambiguous-strand variants (A/T, C/G) and duplicate-position variants. We computed 20 PCs over the merged chromosomes.

Because samples were aggregated into ‘meta-samples’, covariate scores were calculated as the average value of the constituent samples within each meta-sample. For instance, the age of a meta-sample was defined as the average age of all samples included in that group. Similarly, sex was handled by averaging the binary values (0 for female, 1 for male) of the constituent samples.

TCGA validation variants were adjusted for multiple testing using the Benjamini–Yekutieli procedure^36^ (reported q-values), which accounts for the possible hypothesis dependencies expected from germline variants that are in LD.

### P-value Bootstrapping and Adaptive Iteration

The GroupSig framework relies on the random assignment of samples into meta-samples. This stochastic partitioning introduces Monte Carlo variability; a single random partition may, by chance, yield an association p-value that is not representative of the underlying biological signal. To mitigate this and ensure the robustness of our findings, we utilize a bootstrapping approach where the grouping, signature inference, and association testing are repeated across *m* independent random partitions. For each variant-signature pair, we summarize the evidence by calculating the average of the negative log-transformed p-values:

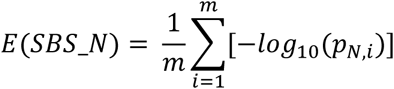

Where SBS_*N* indicates single base substitution signature number N, *P_N,i_* is the p-value reported for the association of SBS_*N*, at iteration *i*. As demonstrated in our calibration analysis (Supplementary methods), averaging evidence across repeated iterations substantially stabilizes the signal and reduces the standard error of the mean (SEM), preventing both spurious false positives and the loss of genuine signals due to “unlucky” partitions.

Performing a high number of iterations (e.g., m=100) for every variant in a genome-wide scan is computationally prohibitive. To handle this burden, we implemented an adaptive iteration scheme that concentrates computational resources on variants showing evidence of a repeatable signal.

The procedure follows an increasingly stringent multi-step filter:

1. Initial Screening: For every variant we performed an initial batch of *m=5* iterations to compute a baseline E_max_ (the maximum evidence across all tested signatures).
2. Adaptive Advancement: If a variant’s E_max_ exceeded a nominal threshold (-log_10_(5×10^-2^)) we perform another batch of iterations on it.
3. Stringent Thresholding: At each subsequent step, the cumulative E_max_ is re-evaluated against a threshold that becomes one order of magnitude stricter (e.g., moving from 5×10^-2^ to 5×10^-3^, and so on).
4. Convergence: This process continues until either the evidence falls below the required threshold for its current batch size or the variant reaches a maximum of 99 iterations.

For visualization, it is necessary to select a single “representative” partition from the bootstrapped set. We define the representative iteration for a specific variant-signature pair as the one whose individual -log_10_(p-value) is closest to the mean evidence E calculated across all successful iterations for that variant. This ensures that the reported effect sizes and visual representations (e.g., regression plots) reflect the central tendency of the bootstrapped evidence rather than a stochastic outlier.

### DFCI panel version harmonization

The DFCI Profile cohort comprises three panel versions that target different gene sets and genomic regions and therefore yield a different typical number of mutations per sample. After filtering, the median number of mutations were 4, 5, and 8 per sample for panels 1, 2, and 3, respectively (Table S3). To harmonize analyses across panels we down sampled mutations from panels 2 and 3 so that their per-sample mutation medians matched the panel 1 median.

Thus we randomly subsampled their mutations by a factor of 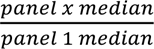 yielding subsampled panels with a median matching the panel 1 median.

### Subsampling TCGA samples

We used the WES data from TCGA to validate GroupSig, generating an emulated panel on which meta-sample aggregation was performed. We compared the inference results between the emulated panel meta-samples and individual WES samples, utilizing the same set of signatures that we applied to DFCI. Emulated panel data is generated by applying similar filters to TCGA as we did to DFCI, then randomly subsampling mutations from each sample to mimic the number of mutations that are observed in targeted panel sequencing.

When both primary and metastatic specimens were available, only the primary tumor was retained; patients with multiple primary tumors were excluded (n = 69). To remove hypermutated outliers we applied per–tumor-type filters: within each tumor type we excluded the top 10% by somatic-mutation count and excluded any sample with >1,000 somatic mutations. After filtering, the median somatic-mutation burden across retained samples was 78 (Table S4).

To emulate panel sequencing, we sampled without replacement each WES sample’s 96-trinucleotide context vector a number of times equal to its subsampling factor. The subsampling factor was chosen so that the post-subsampling median approximated our panel target, while accounting for sample specific mutational burden. We define:

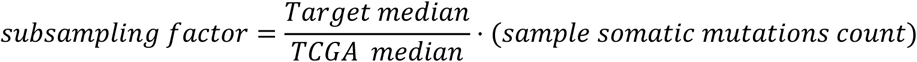

Using as target the DFCI panel 3 median of 8 and TCGA WES median of 78. The product of the factor and the sample mutations count was rounded down.

### DNA Repair Genes Enrichment

To test if sub-threshold SigQTL signals are enriched in relevant biological pathways, we evaluated the enrichment of associated loci within a curated set of 261 high-confidence DNA-repair genes (DRGs) from Gene Ontology^37^ (Supplementary methods). Variants were assigned to genes based on non-redundant gene intervals (5′ UTR to 3′ UTR; refGene hg19^38,39^), excluding those in overlapping intervals of DRGs and non-DRGs to avoid ambiguity. To prevent signal inflation from LD, correlated variants were collapsed into single loci.

To evaluate the biological relevance of associations that did not reach genome-wide significance, we tested whether these sub-threshold putative SigQTL signals preferentially localized to DNA-repair pathways. For each significance threshold *t*, we compared the observed number of SigQTL loci assigned to DRGs (*x*) against the expected number under the null hypothesis that SigQTL loci are uniformly sampled from the background genomic distribution.

This approach assumes that under the null, the proportion of SigQTLs in DRGs should simply mirror the background fraction of total variants located within these genes. Enrichment was formally modeled using a one-sided hypergeometric over-representation test^40^ to determine if the SigQTLs are significantly more frequent within DRGs than would be expected by chance

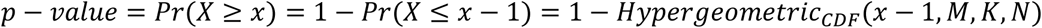

Where *M* is the total background loci, *K* is the number of background loci in DRGs, and *N* is the total number of SigQTL loci at threshold *t*.

## Results

### Age signatures association via GroupSig

We start by evaluating the ability of GroupSig to infer associations between age and age-related signatures. Two signatures, SBS1 and SBS5, are considered canonical ‘clock-like’ age-related processes^34^. SBS1 consists predominantly of C>T substitutions resulting from the spontaneous deamination of 5-methylcytosine^41^, accumulating approximately linearly with age during cell divisions. While the etiology of SBS5 is not fully understood^7,34^, it is associated with chronological time independent of cell division rates. Because both signatures are widely regarded as clock-like, we utilized them to benchmark GroupSig’s ability to recover known biological associations from sparse mutational data.

To establish an age association baseline, we first evaluated the association of age with these signatures in the TCGA WES cohort using the standard individual-sample analysis. To ensure comparability, we applied the same filters used in the GroupSig framework: filtering out samples that are not of European ancestry, excluding the top 10% of samples with the highest mutation count per tumor type (n=6,802) and restricting the analysis to the same set of signatures. Signatures activity was inferred for each sample using MuSiCal. A pan-cancer analysis confirmed that both SBS1 and SBS5 were significantly correlated with age in the full WES dataset (Fig. 2a&b).

**Figure 2.**
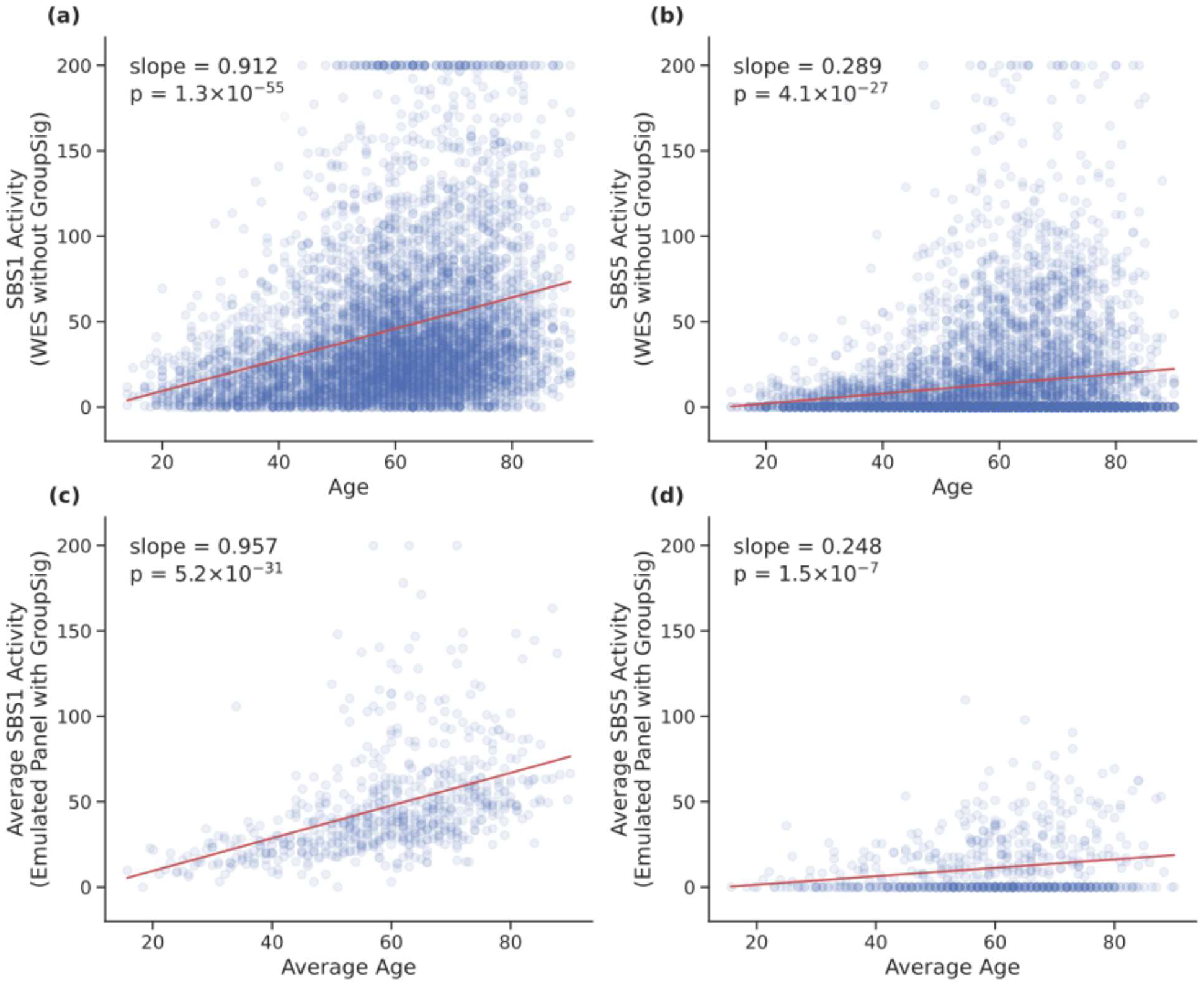
Age association. Linear association between age and mutational signatures activity **(a)** Association between SBS1 activity and age at diagnosis across individual TCGA WES samples (no aggregation) **(b)** Same as **a**, but for SBS5 **(c)** Association between average SBS1 activity and average age at diagnosis across meta-samples aggregated from emulated panel samples using GroupSig **(d)** Same as **c**, but for SBS5. Displayed values are capped at 200.

To emulate the sparsity of panel sequencing in clinical usage, we subsampled mutations in each TCGA sample to match the median mutation counts observed in the DFCI panel 3. This generated a dataset with panel-like sparsity where the ‘ground truth’ of age correlation was already established. We then applied GroupSig to this emulated panel by sorting samples by age and partitioning them sequentially into meta-samples of ten individuals each. Mutational patterns were aggregated within each group, and the meta-sample age was defined as the average age of the constituent samples. Signatures activity were inferred using MuSiCal, followed by correlation analysis which successfully replicated the positive correlation between age and both SBS1 and SBS5 (Fig. 2c&d, Table 1&2). Together, these results demonstrate that GroupSig reliably recovers biologically meaningful age-signature associations from panel-like sparse data, yielding slope estimates within 5% (SBS1) and 14% (SBS5) of the WES benchmarks. This supports applying GroupSig to validate known relationships in sparse clinical sequencing datasets.

**Table 1.**
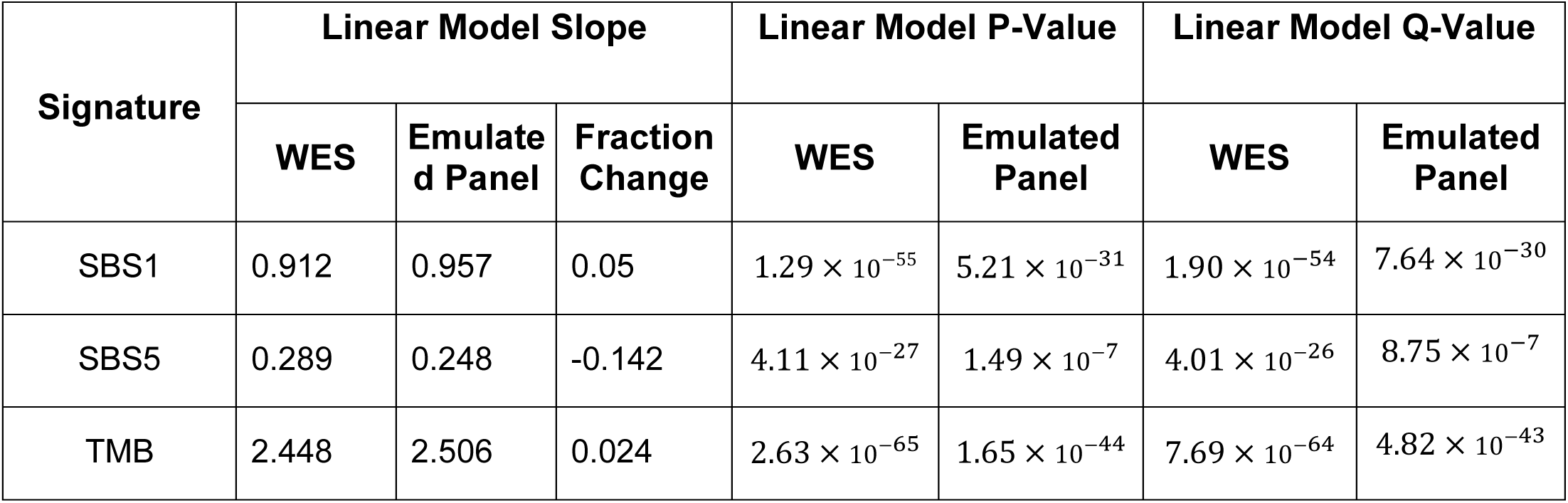
Linear Model.

**Table 2.** Spearman.

| Signature | Spearman Correlation |  | Spearman P-Value |  | Spearman Q-Value |  |
| --- | --- | --- | --- | --- | --- | --- |
|  | WES | Emulated Panel | WES | Emulated Panel | WES | Emulated Panel |
| SBS1 | 0.304 | 0.569 | $9.68 \times 10^{-139}$ | $6.26 \times 10^{-57}$ | $1.42 \times 10^{-137}$ | $1.83 \times 10^{-55}$ |
| SBS5 | 0.072 | 0.137 | $5.85 \times 10^{-9}$ | 0.000476 | $2.14 \times 10^{-8}$ | 0.0002 |
| TMB | 0.36 | 0.527 | $1.26 \times 10^{-197}$ | $9.90 \times 10^{-48}$ | $3.68 \times 10^{-196}$ | $1.45 \times 10^{-46}$ |

### SigQTL - association between Signatures and Genotypes

Having validated GroupSig, we extended the framework to quantify the effect of common germline variants on mutational signatures activity using the DFCI harmonized dataset. We restricted our analysis to a quality-controlled set of 17,076 samples of European ancestry. The final dataset included 3,170,862 variants that passed quality control and were observed in at least 50 samples carrying an alternative genotype (i.e. a minimum of 50 samples with heterozygous genotype or a minimum of 50 samples with homozygous alternative). We should note that the number of samples varies across variants due to imputation constraints.

For each variant, samples were stratified by genotype (homozygous reference, heterozygous, homozygous alternate) and aggregated into meta-samples of size n (*n = 10*). We inferred mutational signatures activity using MuSiCal and assessed the association between each SBS signature activity and the genotype via linear regression controlling for age, sex and the first 20 PCs, which should incorporate sub-populations among Europeans.

We performed genome-wide association analyses across all signatures (Fig. 3a; Fig. S1) and identified 9 SigQTL variants that reached genome-wide significance (p < 5 × 10⁻⁸; Table 3). To ensure robustness to bootstrap-derived variability, we assessed the SEM of −log₁₀(p-value) across 100 bootstrap partitions and required the mean −log₁₀(p-value) to exceed the genome-wide significance threshold by more than 2 SEM. All 9 variants met this criterion (Supplementary Methods; Fig. S2&3).

**Figure 3.**
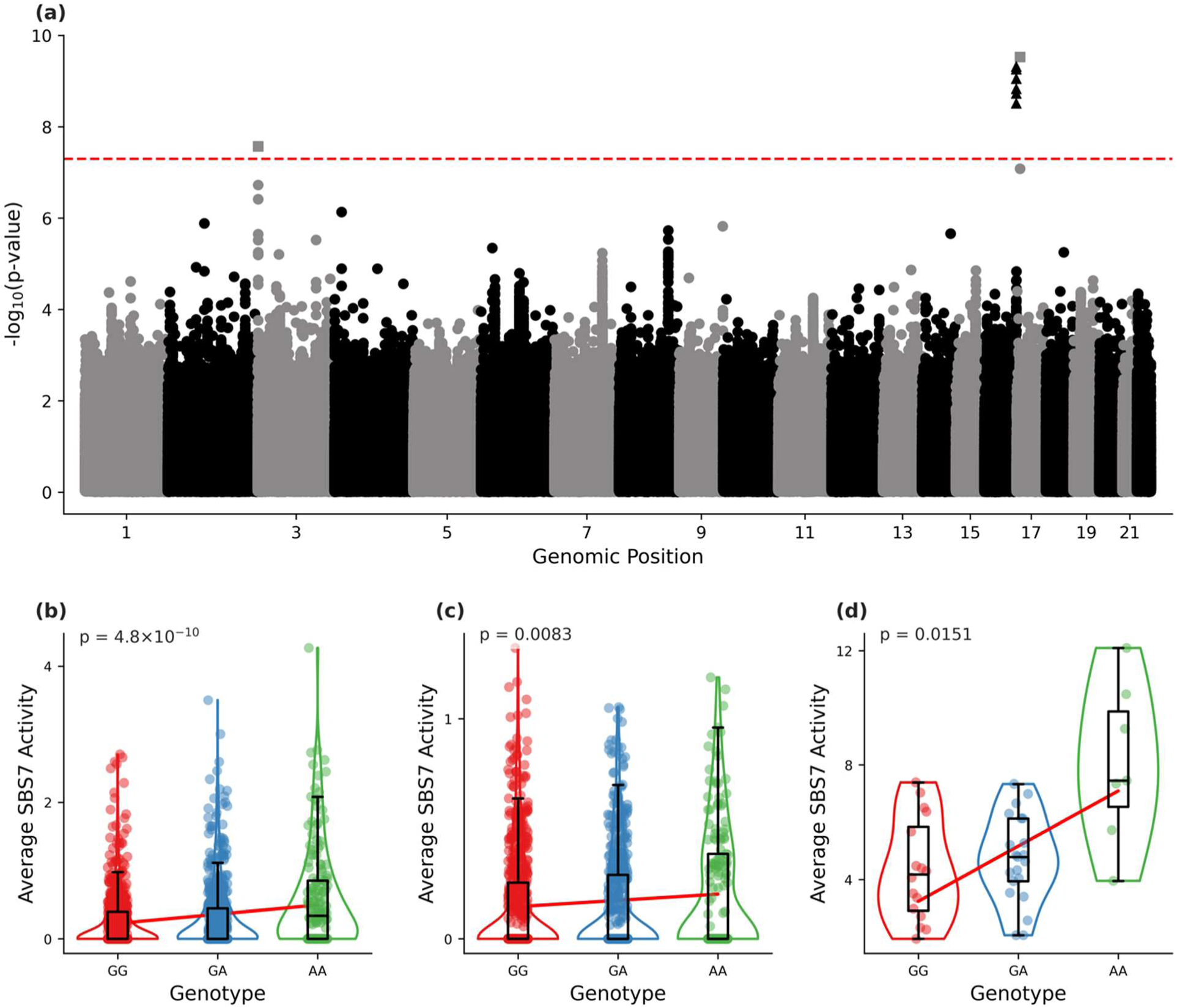
SigQTL results. **(a)** Manhattan plot of the mutational signatures activity GWAS applied to the harmonized DFCI panel. The x-axis represents the variant’s genomic position, the y-axis represents the -log_10_(p-value) of the linear regression models of change in the average SBS7 activity in meta-samples between genotypes at each germline variant. The dotted red line represents the standard GWAS threshold − log_10_(5 × 10^-8^). SigQTLs in locus 16q24.3 are represented as triangles, rs17883687 located at locus 17p13.1 is represented as a square. (**b-d**) Linear association between the rs12448464 variant and meta-samples average SBS7 activities in the DFCI harmonized panel, showing GroupSig results for **(b)** all samples, **(c)** all samples excluding melanoma, and **(d)** melanoma samples only. Genotypes are encoded GG = 0 (homozygous major), GA = 1 (heterozygous), AA = 2 (homozygous minor). Each point is a meta-sample (group).

**Table 3.**
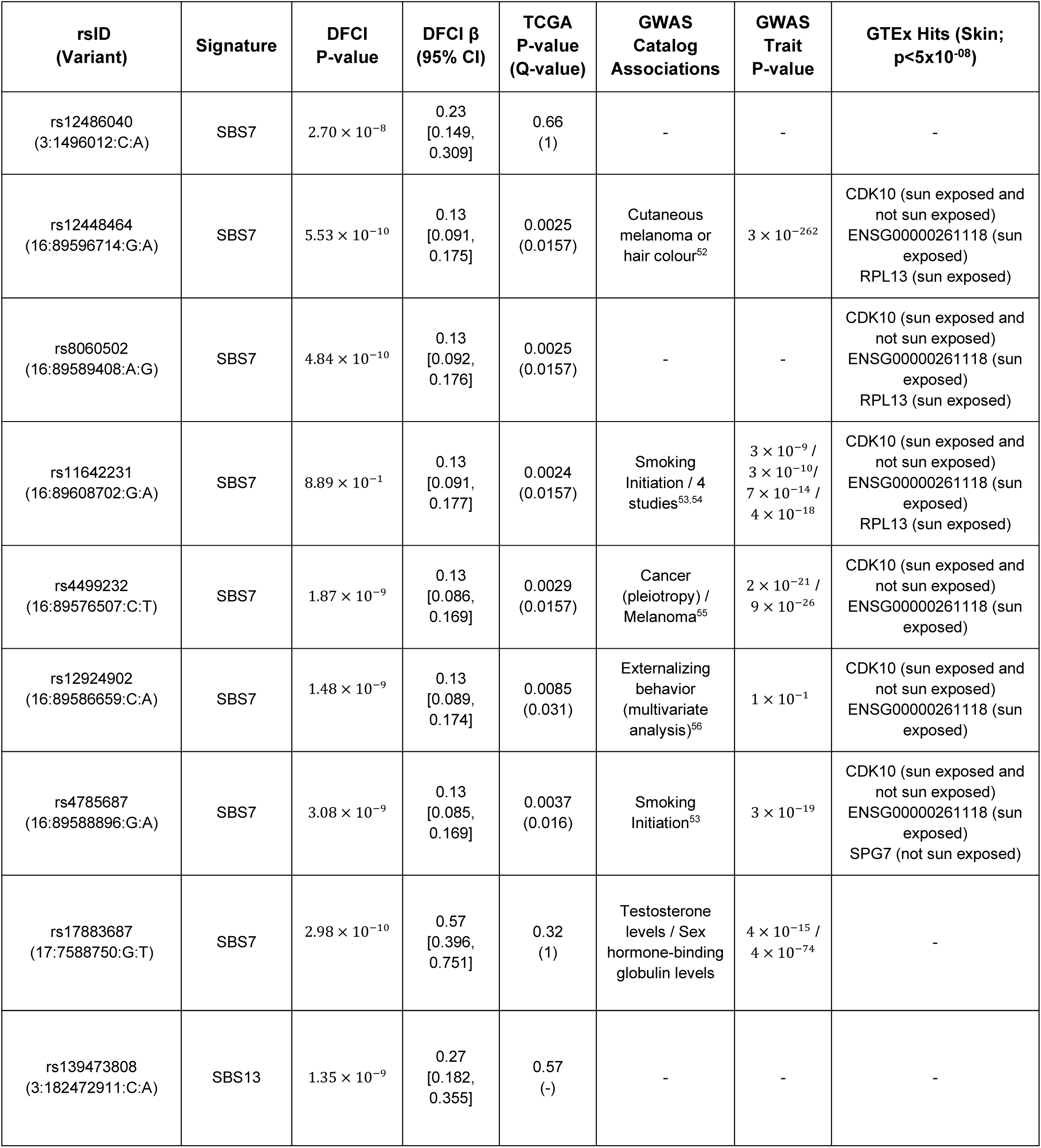
SigQTL variants summary.

The SigQTLs included eight variants associated with SBS7 (UV exposure) and one variant associated with SBS13 (APOBEC); however, this was not the variant previously reported to be associated with APOBEC activity^8^. The most prominent signal mapped to the 16q24.3 locus, where six variants were associated with increased SBS7 activity. The effect sizes of these variants were similar, corresponding to ∼0.13 [0.125-0.132] additional SBS7-attributed mutations per sample in the DFCI panel 1, which extrapolates to ∼6 additional mutations per whole-exome and ∼300 per whole-genome sequencing sample. Four of these variants (rs12448464, rs4499232, rs11642231, and rs4785687) have previously been associated with pigmentation, risk of melanoma and other cancer types, or smoking-related traits (Table 3).

Because SBS7 is highly enriched in melanoma, we tested whether the pan-cancer SBS7 association was driven by chance melanoma-case stratification at these loci. We reanalyzed SBS7 associations in two ways: (i) restricting to melanoma tumors only and (ii) excluding melanoma from the pan-cancer cohort. Of the seven SBS7 SigQTLs identified in the pan-cancer analysis, five remained significant in both the melanoma-only and non-melanoma subsets (Fig. 3b&c&d; Fig. S4). The persistence of these associations after separating melanoma cases reduces the likelihood that the pan-cancer SBS7 signal is explained solely by melanoma enrichment.

Additionally, we identified a variant rs17883687, located in a *TP53* intron at 17p13.1 to be associated with SBS7. This variant has previously been linked to sex-hormone binding globulin and testosterone levels (*p* ≤ 4 × 10^-14^ and 4 × 10^-74^ respectively) - phenotypes implicated in cancer susceptibility^42–45^. The remaining SigQTLs rs12486040 at 3p26.3, and rs139473808 were not previously associated with any phenotype, in GWAS studies.

We further assessed the biological context of our findings by examining LD between SigQTLs and known cancer susceptibility loci, using their R^2^ value (the squared correlation between allelic dosages). Querying the GWAS Catalog for proxy variants (R^2^> 0.6) with reported phenotype associations, we found that the SBS7-associated variant rs17883687 (TP53 intron) is in perfect LD (R^2^= 1) with rs17880847 in the Iberian (Spain) and Utah European reference panels, and in moderate LD in the British panel (R^2^ = 0.65). Rs17880847 has been previously associated with inherited cancer-predisposition syndromes^46–48^.

### SigQTL Validation

To validate the SigQTLs identified in the DFCI targeted panel cohort, we analyzed the TCGA WES dataset. This validation serves a dual purpose: it tests the replicability of the biological associations in a distinct cohort and, crucially, verifies that the findings are not artifacts of the GroupSig aggregation procedure. Unlike targeted panels, WES data provides large enough mutational data to allow for reliable mutational signature inference at the individual sample level, thereby obviating the need for sample grouping.

To ensure strict comparability, we processed the WES data using the same steps we applied to the DFCI cohort: (i) Including only samples of European ancestry (ii) excluding hypermutated samples with high mutation count (top 10% of each tumor type) (iii) using the same signature set (iv) and using MuSiCal for signature inference. We then assessed the association between the SigQTLs that we found and per-sample signatures activity using a linear regression model.

Of the 9 significant variants we found in the DFCI dataset, six SigQTLs were significant after correcting for multiple testing (q<0.05; BY) and present the same trend as in the DFCI dataset (Table 3, Fig. 4a). Of note, the three remaining variants (rs17883687, rs12924902 and rs139473808), while not significant, still showed a trend for higher associated signature activity in the minor variant (Table S5). All of the 6 validated SigQTLs are on the same 16q24.3 locus.

**Figure 4.**
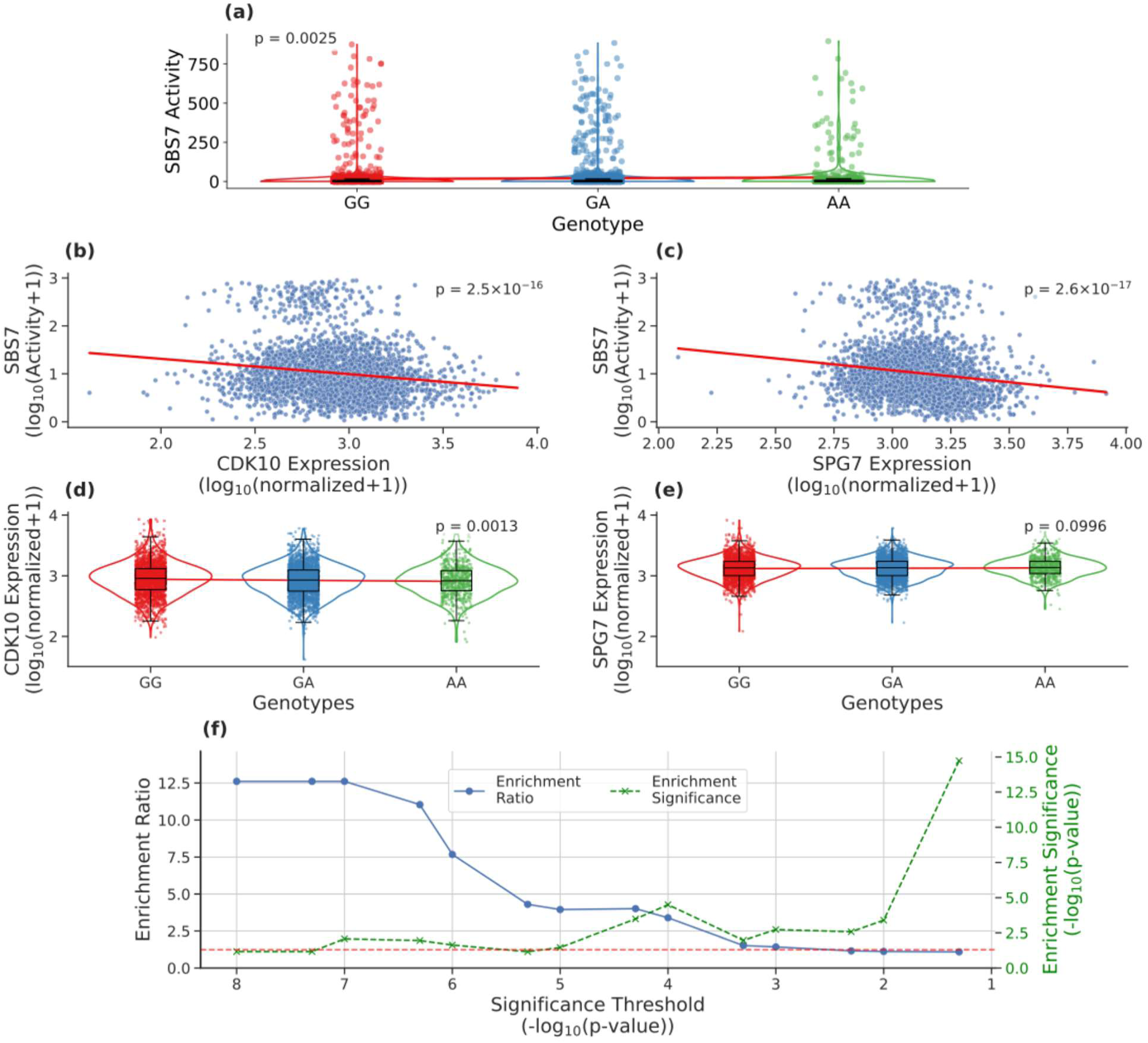
Validation and enrichment results. **(a)** Linear association between the rs12448464 variant and SBS7 activity. Each point is one TCGA WES sample for which mutational signatures were inferred at the sample level (no aggregation). (**b-c**) Linear association between gene expression and SBS7 activity. Each point is one TCGA WES sample for which mutational signatures were inferred at the sample level (no aggregation) **(b)** CDK10 **(c)** SPG7 (**d-e**) Linear association between the rs12448464 variant and gene expression. Each point is one TCGA WES sample **(d)** CDK10 **(e)** SPG7 **(f)** Enrichment analysis over all signatures. The left y-axis shows Enrichment fold (ratio), The right y-axis shows enrichment significance(-log_10_(p-value)), x-axis shows association significance thresholds in -log_10_(p-value). The blue curve represents the enrichment ratio observed for each significance threshold, the green curve represents the significance of the enrichment. The red dotted line (y=-log_10_(0.05)) represents the standard null-hypothesis rejection threshold that the enrichment significance needs to surpass to be considered as significant.

To investigate the biological mechanisms underlying these signals, we cross-referenced our validated SigQTLs with the GTEx database^13^. All the variants at the 16q24.3 locus were found to be expression quantitative trait loci (eQTLs) for the CDK10 oncogene^49^ in both sun-exposed and non-exposed skin tissues. Additionally, four of these variants (Table 3) were associated with the expression of *SPG7* in skin.

To assess whether *CDK10* and *SPG7* are linked to SBS7 activity, we analyzed the TCGA cohort and tested for associations between gene expression and SBS7 activity. In TCGA, the RNA expression levels of both *CDK10* and *SPG7* were negatively correlated with SBS7 activity (Fig. 4b&c). In addition, the SBS7 minor allele at the 16q24.3 locus was associated with lower *CDK10* RNA expression (Fig. 4d). The lower levels of *CDK10* were consistent with the association between this locus and increased SBS7 activity. In contrast, there was no significant correlation between genotypes and *SPG7* expression (Fig. 4e), suggesting a stronger connection between the 16q24.3 locus and *CDK10* than *SPG7*.

Although these genes are not members of the canonical DNA repair machinery, their association with SBS7 activity suggests that the “mutational phenotype” is influenced by broader cellular homeostatic processes. For instance, *CDK10* is a known regulator of the G2/M transition; its modulation may dictate the temporal window available for nucleotide excision repair to process UV lesions before they are fixed as mutations during replication. Alternatively, *SPG7*’s role in mitochondrial proteostasis and the maintenance of the mitochondrial permeability transition pore could influence the accumulation of oxidative stress or alter the energy-dependent kinetics of DNA repair pathways. These findings imply that the 16q24.3 locus may modulate UV-signature activity by altering the cellular context in which DNA damage is sensed and processed.

### DNA Repair Genes are Enriched in Sub-threshold SigQTLs

Our SigQTL analysis primarily identified common variants with large effects; e.g. the variants in the 16q24.3 locus have an increase of ∼0.13 [0.125-0.132] (Table 3) mutations per allele in DFCI. This is an increase of ∼40% compared to the baseline signature activity of 0.303 [0.249-0.317] (i.e. for the major allele Table S6). However, given the expected polygenic nature of DNA maintenance, many variants likely exert subtler influences (∼2%–10%) that fall below the stringent genome-wide significance threshold.

To quantify the extent of this latent signal, we compared the frequency of SigQTLs within DRGs to their frequency across all genes. Although variants within DRGs constitute only 1.13% of the tested variants (N = 1,278,773), we observed an overrepresentation of DRGs among our association hits. Under the null hypothesis, the proportion of SigQTLs mapping to DRGs should mirror the genomic baseline; however, our results demonstrate a significant enrichment that scales with statistical stringency.

Specifically, we found a 12.6-fold enrichment for DRGs at the genome-wide significance threshold (p < 5×10^-8^), which gradually decayed to a 1.3-fold enrichment as the threshold was relaxed to a nominal level (p = 0.05; Fig. 4f). This robust enrichment gradient suggests that while many individual variants lack the power to surpass a stringent GWAS cutoff, the “suggestive” signal space is systematically anchored in the biological machinery of DNA maintenance.

## Discussion

The massive scale of clinical targeted panels now represents the dominant source of tumor sequencing data, routinely achieving cohort sizes that are unattainable for whole-exome or whole-genome studies. However, despite this scale, panel cohorts have remained underutilized for mechanistic studies of mutational processes because individual tumors typically harbor too few mutations for stable signature inference. We addressed this “sparsity barrier” by introducing GroupSig, an aggregation framework that treats inter-individual variability in mutational processes as a measurable quantitative trait. By shifting the unit of inference from the fragmentary individual sample to the population stratum, GroupSig turns cohort scale into a tool for signal denoising. Applying this framework to a cohort of 31,984 samples and validating key signals in an independent dataset, we provide direct evidence that common germline variation contributes to systematic differences in the activity of somatic mutational processes.

The strongest and most reproducible signal identified in our scan mapped to 16q24.3 and was associated with increased SBS7 activity. As the canonical UV-associated signature, the association of SBS7 with inherited variation suggests a genetically encoded modulation of exposure sensitivity; importantly, the persistence of this signal outside melanoma cases argues that the effect is a genuine biological modifier rather than a case-enrichment artifact. The discovery that the 16q24.3 locus involves *CDK10* and *SPG7*, genes outside the canonical DNA repair machinery, suggests that mutational variability may be influenced by broader cellular contexts beyond the core “repair-ome”. We hypothesize that these genes might act as “state-setting” modifiers. For instance, *CDK10* regulation of the G2/M transition could potentially modulate the temporal window available for lesion repair before replication. Alternatively, *SPG7* might indirectly influence repair capacity through its roles in mitochondrial proteostasis and metabolic state. In this view, inherited variation may tune the cellular environment, affecting variables like replication timing or stress signaling, to shape somatic mutational outcomes.

Beyond individual lead loci, the systematic enrichment of sub-threshold signals in DNA-repair genes supports a truly polygenic model for mutational process regulation. The overrepresentation of DNA-repair genes among suggestive associations implies that while only a small number of loci exceed genome-wide significance at current sample sizes, the “gray zone” of the Manhattan plot is systematically anchored in the biological machinery of DNA maintenance. This motivates the development of a “Polygenic Mutational Score,” which could eventually predict an individual’s intrinsic sensitivity to specific mutational processes.

Despite these advances, several limitations remain. Pan-cancer aggregation may dilute tissue-specific SigQTLs, and signatures associated with environmental exposures remain vulnerable to residual confounding by anatomical site or behavioral proxies. Furthermore, current signature reference sets and ancestry restrictions limit the generalizability of these findings across diverse populations. In addition, GroupSig produces group-level activity estimates that depend on meta-sample construction and filtering choices, and these choices can influence apparent effect sizes. Finally, like in any GWAS analysis, statistical association at non-coding loci requires follow-up to resolve the causal variants, target genes, and mechanisms linking germline variation to signature activity.

From a statistical perspective, GroupSig departs from the conventional “high-N” dogma, which typically favors many low-reliability observations over fewer high-fidelity ones. This pivot is necessitated by the “information floor” inherent to mutational signature deconvolution; because signature inference requires a minimum mutational density to reach mathematical stability, individual sparse samples often fall below the threshold required for robust inference. In this non-standard regime, aggregating individual samples into high-fidelity meta-samples is not merely a smoothing technique but a fundamental requirement for enabling measurement. Consequently, while individual-level analysis remains the standard for traditional GWAS where the phenotype is directly observable, the inherent sparsity of panel data requires the GroupSig framework to reconstruct the latent phenotype at the population level.

The utility of the GroupSig framework extends beyond SigQTL discovery to any research question involving sparse mutational signatures data. For instance, it can be deployed to test whether specific signature activities are enriched in clinical subgroups, such as treatment responders versus non-responders, or to quantify the impact of environmental carcinogen exposures across both discrete and continuous variables.

However, it is essential to distinguish GroupSig’s role as a discovery-oriented population tool from individual-level inference methods. While tools like SigMA^50^ or Mix^51^ were specifically engineered to deconvolve signatures in a single sparse sample for diagnostic purposes, GroupSig is optimized for identifying broader biological and genetic determinants. By shifting the unit of inference to the population stratum, it reveals systematic associations that remain obscured by stochastic noise at the level of the individual patient.

Integrating these germline determinants into precision oncology will enable us to move beyond predicting cancer risk to predicting how a tumor will accumulate mutations, providing the foresight necessary to tailor surveillance and therapy to a patient’s unique genetic background.

## Data Availability

TCGA data are publicly available from the Genomic Data Commons (https://portal.gdc.cancer.gov/)

DFCI PROFILE data are protected due to patient privacy. All summary statistics are available within the supplementary tables.

Analysis code and pipelines are available at GitHub: https://github.com/alonravid1/SigQTL

## Acknowledgments

This work was supported by the Israel Science Foundation (ISF), grant number 2794/21.

