## Supplemental Methods and Figures for "Breaking the sparsity barrier in clinical targeted-panel sequencing: Mapping the inherited determinants of mutational signatures"

### Supplementary Methods

#### DNA Repair Genes Enrichment

To test if sub-threshold SigQTL signals are enriched in relevant biological pathways, we evaluated the enrichment of associated loci within DNA-repair genes (DRGs) from Gene Ontology. To do so we: (i) retrieved a list of high-confidence DRGs. (ii) assigned variants to one of three categories (DRGs, non-DRGs, not in genes) based on their genomic span (5′UTR to 3′UTR). (iii) excluded variants in overlap between DRGs and non-DRGs to avoid ambiguous assignments. (iv) collapsed correlated SigQTLs into a single locus to avoid signal inflation from linkage disequilibrium (LD).

We obtained DRGs from Gene Ontology by selecting the "DNA repair" term in their genes-annotation and restricting to human genes supported by direct experimental evidence for DNA-repair function, yielding 261 genes. GO-provided gene synonyms were retained to maximize mapping sensitivity.

Gene coordinates were derived from the hg19 refGene.gtf annotation. For each gene we used the minimal 5′UTR start and the maximal 3′UTR end across annotated transcripts to define a non-redundant genomic span. Variants were categorized as: DRG (falls within a DRG span), non-DRG (falls within a non-DRG gene span), or not-in-gene (falls outside any gene span, excluded from the analysis). Variants overlapping two or more gene spans of different categories were labelled as "overlap" and were excluded from enrichment testing (approximately 8% of variants).

Because LD introduces local correlation among germline variants and violates the independence assumption of the hypergeometric test, we collapsed correlated SigQTLs into loci using a single-linkage procedure performed separately for each chromosome and each mutational signature. Starting from the most 5′ variant, consecutive SigQTLs were merged into the same locus if the distance between adjacent variants was ≤500 kilobase pairs (Kbps); the locus was extended while subsequent adjacent variants remained within 500 Kbps. Single variants with no neighbor within 500 Kbps were also treated as loci. Each finalized locus was localized to the SigQTL with the strongest signal (defined as smallest p-value), and the locus was then classified as DRG, non-DRG, or overlap according to that representative SigQTL.

#### Calibration and power

##### Pilot set and extrapolation

Before applying GroupSig to test associations between germline variants and mutational signatures activity, we quantified how the stochastic partitioning step affects the resulting association p-values. Because samples are randomly partitioned into meta-samples, a single partition can, by chance, yield an unusually strong or weak association, creating both false positives and false negatives. To reduce this Monte Carlo variability, we repeated the grouping step multiple times per variant and summarized evidence using the average of the resulting $-\log_{10}$(p-values).

We calibrated this variability in a pilot experiment on ~10,000 randomly selected variants. For each variant and each signature tested, we: (i) stratified samples by genotype and randomly aggregated them into meta-samples (ii) inferred signatures activity for each meta-sample using MuSiCal (iii) tested the association between genotype and inferred signature activity. We repeated steps (i)–(iii) for 100 independent random partitions for each variant.

For each variant–signature pair we summarized evidence by taking the average of $-\log_{10}$(p-value) across 100 independent random partitions and calculated the resulting standard deviation (SD) of the $-\log_{10}$(p-values). In our pilot set the SD increased with signal strength, reaching ~0.8 for highly significant associations (Fig. S2).

Because the pilot set did not contain strong signals, these observations did not directly inform variability at larger $-\log_{10}$(p-values); consequently, we extrapolated the empirical relationship between mean $-\log_{10}$(p-values) and its SD to cover stronger hypothetical signals. For extrapolation we fitted three simple functional forms to the pilot data: linear, power-law and logarithmic. We expect the data to follow a power-law, which we used for the calibration, and the linear and logarithmic later on to compare with the power-law extrapolation on a validation set.

To estimate the power-law parameters, we begin with its formulation:

$$y = a \cdot x^b$$

By applying log to both sides:

$$log_{10}(y) = log_{10}(a \cdot x^b) = log_{10}(a) + log_{10}(x^b) = log_{10}(a) + b \cdot log_{10}(x)$$

We can rewrite this as a linear regression with variables $y' = log_{10}(y), x' = log_{10}(x)$

$$y' = log_{10}(a) + b \cdot x' = a' + b \cdot x'$$

We can calculate the intercept $a'$ and slope $b$, calculate $a$ by substituting:

$$a = 10^{log_{10}(a)} = 10^{a'}$$

Using SBS7 estimates and the power-law for which the best fit was observed ($a = 0.37$, $b = 0.54$), we get for some hypothetical SigQTL with an average $-\log_{10}(3 \times 10^{-8}) = 7.52$ (which passes the genome-wide significance threshold of $-\log_{10}(5 \times 10^{-8}) = 7.3$ by a modest amount) we extrapolate a SD of:

$$SD = 0.37 \cdot 7.52^{0.54} \approx 1.1$$

With 100 independent partitions the SEM is calculated as such:

$$SEM = \frac{SD}{\sqrt{iterations}} = \frac{1.1}{\sqrt{100}} = 0.11$$

And the 95% lower confidence bound for the average evidence is calculated as such:

$$CI_{95\%} = average\ - z_{95\%} \cdot SEM = 7.52\ - 1.96 \cdot 0.11 = 7.304$$

Only associations with true evidence very close to the genome-wide significance cutoff $-\log_{10}(5\times10^{-8}) = 7.3$ are likely to change classification because of partitioning noise, while associations whose true −log10(p-value) lies well above this range are unlikely to be missed. Null variants in the pilot set failed to surpass an upper average −log10(p-value) of 5. Using a conservative upper limit of 6, we get the following upper confidence interval bound for 95% CI:

$$CI_{95\%} = 6\ + 1.96 \cdot 0.11 = 6.2156$$

Associations that are well below the GWAS threshold therefore unlikely to be spuriously called SigQTLs. These results show that averaging $-\log_{10}$(p-value) across repeated partitions substantially mitigates false positives and false negatives induced by stochastic grouping.

#### Batch size calibration

We then calibrated the number of iterations performed in each batch of our adaptive iteration scheme. Ideally, we would simply apply 100 iterations for each variant in our dataset to produce robust results. However, each additional iteration produces a new independent partition of samples which requires an additional mutational signatures inference, followed by linear-regression fitting. These operations dominate the cost of our pipeline (~15s per iteration on our machines) and are many orders of magnitude more resource consuming than any other repetitive step. Running SigQTL over ~3,100,000 variants will therefore take ~1,300,000 hours (even using ~300 CPUs in parallel the analysis will take ~4,800 hours).

To reduce computational cost while preserving sensitivity, we implemented a two-step, iteration-based workflow. Each variant is first processed in an initial screening batch of *m* GroupSig iterations; only variants that pass this first screen are processed in subsequent batches. Under the null hypothesis, p-values are approximately uniformly distributed on (0,1), meaning an initial screening with a threshold of 1.3 ($-\log_{10}(0.05)$) will exclude roughly 95% of null variants in the first pass, substantially shrinking the set of candidate SigQTL variants. The batch size *m* controls the precision of the screening estimate: we choose *m* to ensure that GroupSig has high sensitivity in the initial screen, thereby avoiding the inadvertent removal of true SigQTLs. This two-stage design concentrates expensive, full-resolution bootstrapping on a much smaller subset of promising variants and makes the overall workflow computationally feasible.

Using *m = 2* iterations in the screening batch yields a 99.99% confidence-interval lower bound for true SigQTLs of:

$$CI_{99.99\%} = 7.3 - 3.89 \cdot \frac{1.1}{\sqrt{2}} = 4.275$$

Showing that even under stringent confidence requirements, a SigQTL's average $-\log_{10}$(p-value) is expected to stand well above the initial screening threshold. To guard against departures from normality while maintaining feasibility, we conservatively used *m = 5* iterations.

#### Extrapolation validation

To validate our preliminary calibration, we compared the extrapolation functions fitted on the pilot set on a test set of putative associations derived from the GWAS results (average −log10(p-value) > 5; Fig. S3). The power-law model produced the lowest root-mean-square error (RMSE) across signatures, with one exception: for SBS1 the logarithmic fit performed marginally better (RMSE improvement ~ 4%).

### Supplemental Figures

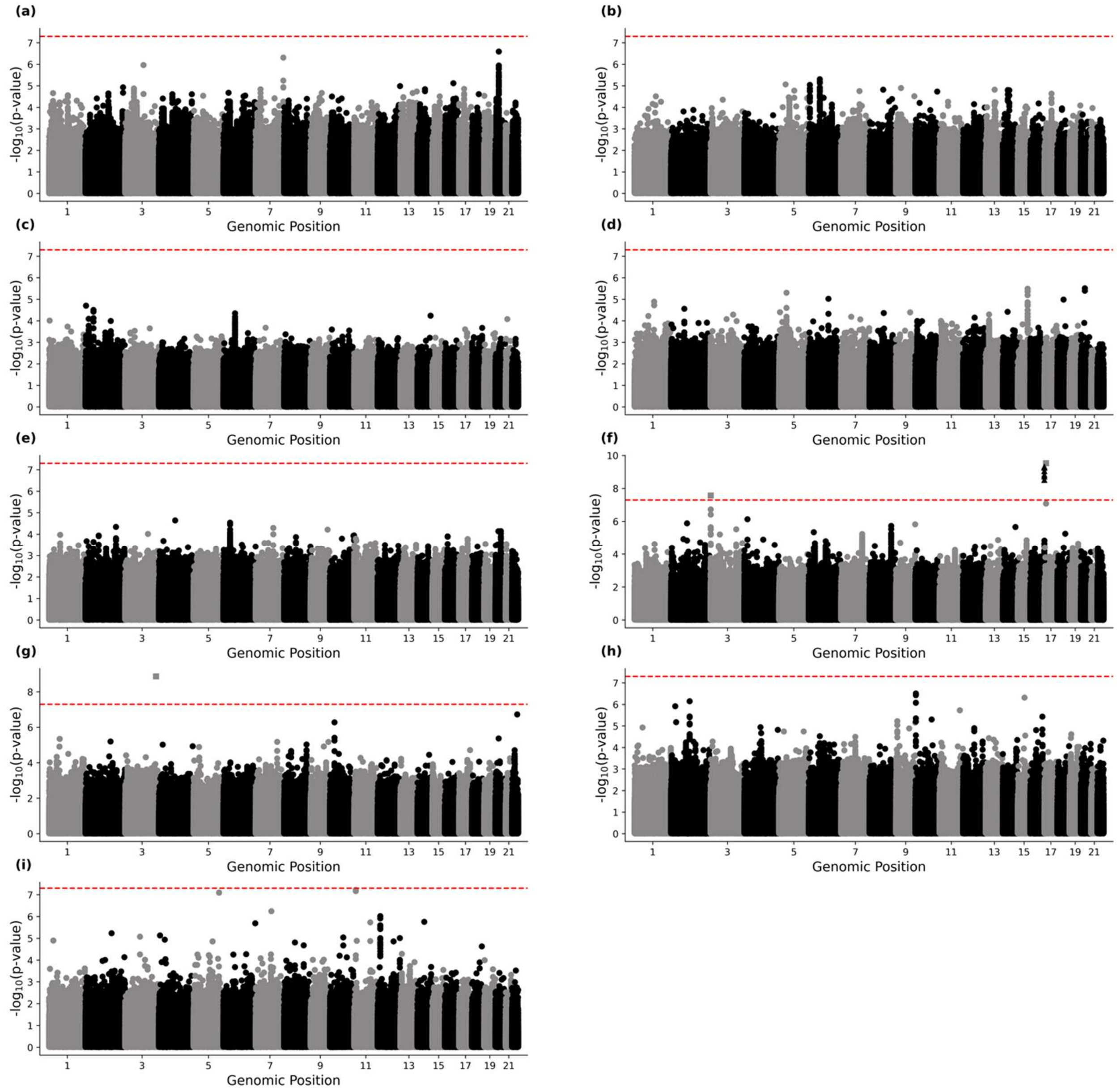


Figure S1 - Manhattan plots of germline variants association with signature activity across the genome: **(a)** SBS1 **(b)** SBS2 **(c)** SBS3 **(d)** SBS4 **(e)** SBS5 **(f)** SBS7 **(g)** SBS13 **(h)** SBS17 **(i)** SBS18

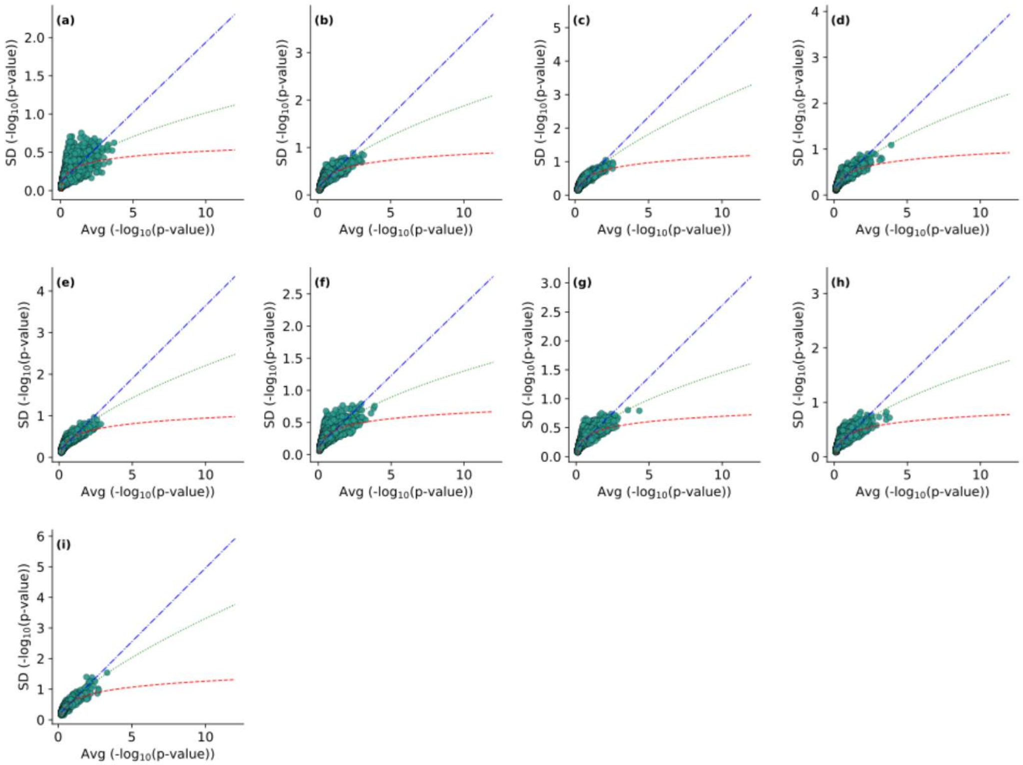


Figure S2 - The observed and extrapolated relations between average log-transformed significance level (-log$_{10}$(p-value)) and standard deviation of log-transformed significance (-log$_{10}$(p-value)) per-variant, resulting from 100 independent GroupSig iterations. Extrapolation was performed using linear (blue curve), power-law (green curve) and logarithmic (red curve) extrapolations. Each point is the result of testing the association between a variant in the pilot set (green) for **(a)** SBS1 **(b)** SBS2 **(c)** SBS3 **(d)** SBS4 **(e)** SBS5 **(f)** SBS7 **(g)** SBS13 **(h)** SBS17 **(i)** SBS18

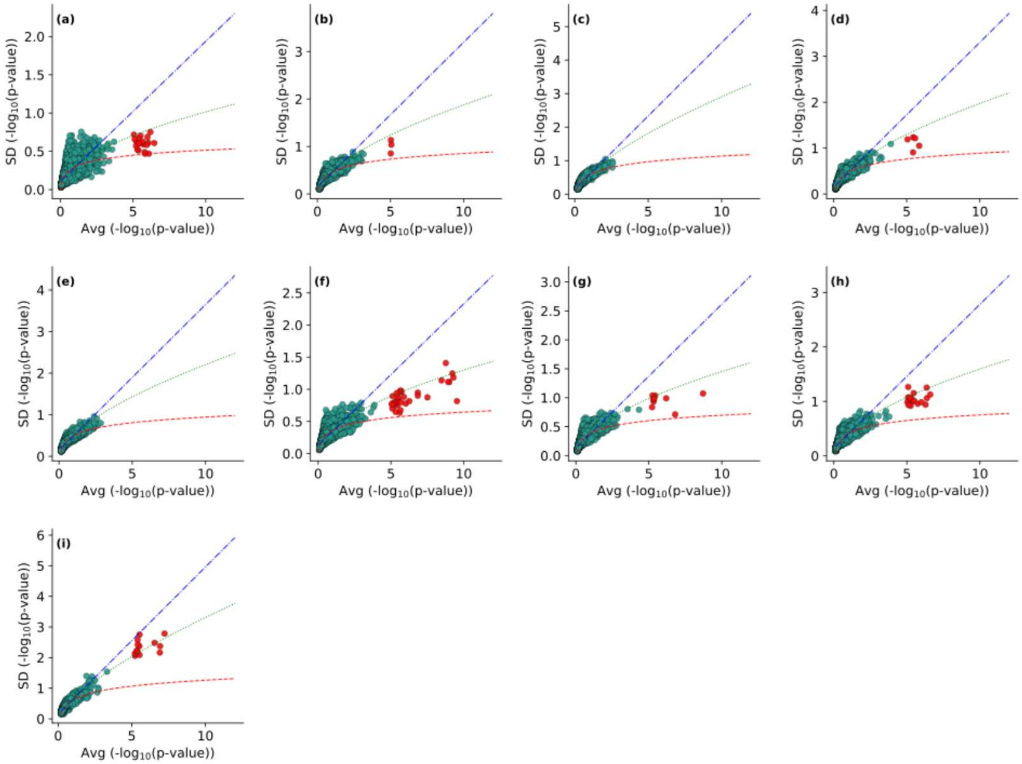


Figure S3 - The observed and extrapolated relations between average log-transformed significance level ($-\log_{10}$(p-value)) and standard deviation of log-transformed significance ($-\log_{10}$(p-value)) per-variant, resulting from 100 independent GroupSig iterations. Extrapolation was performed using linear (blue curve), power-law (green curve) and logarithmic (red curve) extrapolations. Each point is the result of testing the association between a variant in the pilot set (green) or putative SigQTLs($p<10^{-5}$ in at least one signature association; red) for **(a)** SBS1 **(b)** SBS2 **(c)** SBS3 **(d)** SBS4 **(e)** SBS5 **(f)** SBS7 **(g)** SBS13 **(h)** SBS17 **(i)** SBS18

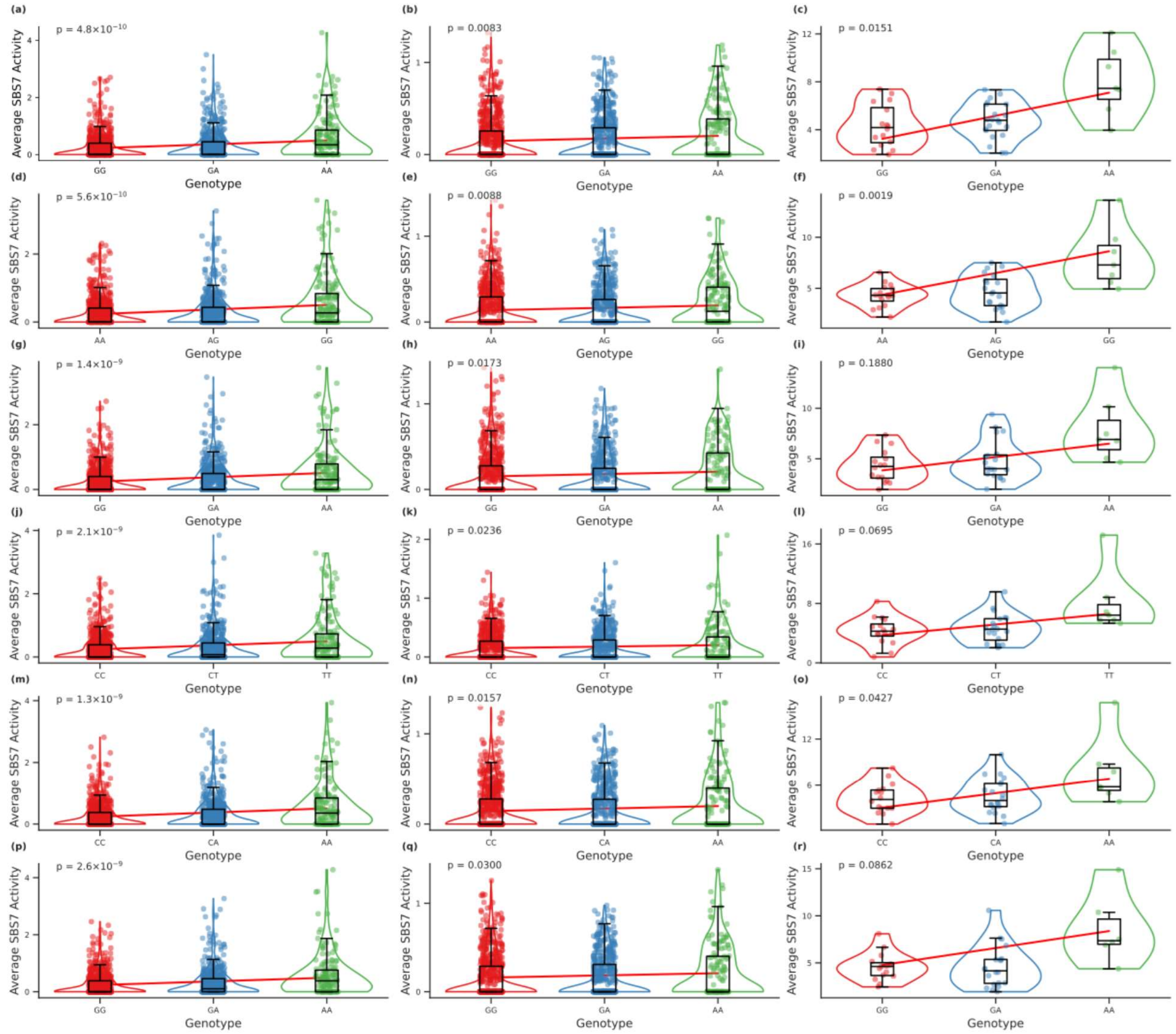


Figure S4 - Variant-level SigQTL plots for SBS7 loci with adequate sample size. For each locus, three figure panels compare SBS7 activity across genotype groups in different cohorts: all samples (first panel), excluding melanoma samples (middle panel), and melanoma samples only (last panel). Panels **a–c**: variant 16:89596714:G:A; **d–f**: variant 16:89589408:A:G; **g–i**: variant 16:89608702:G:A; **j–l**: variant 16:89576507:C:T; **m–o**: variant 16:89586659:C:A; **p–r**: variant 16:89588896:G:A. Each plot displays SBS7 signature activity stratified by genotype (x-axis) with the corresponding group distributions and statistical annotations.
